# *De novo* genome assemblies from two Indigenous Americans from Arizona identify new polymorphisms in non-reference sequences

**DOI:** 10.1101/2023.10.23.563520

**Authors:** Çiğdem Köroğlu, Peng Chen, Michael Traurig, Serdar Altok, Clifton Bogardus, Leslie J Baier

## Abstract

There is a collective push to diversify human genetic studies by including underrepresented populations. However, analyzing DNA sequence reads involves the initial step of aligning the reads to the GRCh38/hg38 reference genome which is inadequate for non-European ancestries. To help address this issue, we created a modified hg38 reference map using *de novo* sequence assemblies from Indigenous Americans living in Arizona (IAZ). Using HiFi SMRT long-read sequencing technology, we generated *de novo* genome assemblies for one female and one male IAZ individual. Each assembly included ∼17 Mb of DNA sequence not present (non-reference sequence; NRS) in hg38, which consists mostly of repeat elements. Forty NRSs totaling 240 kb were uniquely anchored to the hg38 primary assembly generating a modified hg38-NRS reference genome. DNA sequence alignment and variant calling were then conducted with WGS sequencing data from 387 IAZ cohorts using both the hg38 and modified hg38-NRS reference maps. Variant calling with the hg38-NRS map identified ∼50,000 single nucleotide variants present in at least 5% of the WGS samples which were not detected with the hg38 reference map. We also directly assessed the NRSs positioned within genes. Seventeen NRSs anchored to regions including an identical 187 bp NRS found in both de novo assemblies. The NRS is located in *HCN2* 79 bp downstream of exon 3 and contains several putative transcriptional regulatory elements. Genotyping of the *HCN2*-NRS revealed that the insertion is enriched in IAZ (MAF = 0.45) compared to Caucasians (MAF = 0.15) and African Americans (MAF = 0.03). This study shows that inclusion of population-specific NRSs can dramatically change the variant profile in an under-represented ethnic groups and thereby lead to the discovery of previously missed common variations.

**AUTHOR SUMMARY:** GRCh38/hg38 reference genome has been the standard reference for large-scale human genetics studies. However, it does not adequately represent sequences of non-European ancestry. In this study, using long-read sequencing technology, we constructed *de novo* sequence assemblies from two Indigenous Americans from Arizona. We then compared the *de novo* assemblies to the hg38 reference genome to identify non-reference sequences (NRSs). We integrated these NRSs into our whole-genome sequencing (WGS) variant calling pipeline to improve read alignment and variant detection. We also directly assessed the NRSs positioned within genes. Inclusion of population-specific NRSs dramatically changed the variant profile of our study group with under-represented ethnicity, revealing common variation not detected by our previous population-level WGS and genotyping studies.

## INTRODUCTION

The lack of diversity in human genetic studies is of growing concern, particularly as personalized medicine is becoming a reality. The majority of genomics data, approximately 80 percent, predominantly represents individuals of European ancestry, highlighting a significant underrepresentation of other populations [1]. This lack of diversity has had adverse consequences in genetic testing and variant interpretation, particularly for patients of African and Asian ancestry [2,3]. Recent large-scale genome studies, such as the All of Us initiative [4], the Human Heredity and Health in Africa (H3Africa) project [5], and the GenomeAsia100K project [6], have made significant efforts to address this disparity by actively increasing the representation of diverse populations. However, aside from the challenges associated with sample collection for sequencing, these endeavors face two main limitations when generating genomic data for underrepresented populations. One limitation is due to short-read sequencing being the main method in large population studies. While short-reads offers valuable insights through comparisons of allele frequencies and functional annotations, they struggle to capture rare variants and accurately characterize structural variants in isolated populations [7–9]. This restricts the comprehensive understanding of genetic diversity and hampers the identification of variants associated with diseases. The second limitation stems from the reliance on a human genome reference during DNA sequence read analysis. The current reference map, the GRCh38/hg38 reference genome, primarily comprises sequences derived from a single DNA donor of African-European origin, accounting for approximately 70% of the primary assembly sequences [10]. Consequently, the current reference map still exhibits limitations in adequately representing the genetic diversity found within diverse populations. It is important to recognize that individuals may carry unique sequences that are not represented in the reference map, thereby harboring genomic regions with potential disease associations and other phenotypic relevance. Therefore, it has been imperative for the genomic research in isolated populations to move beyond the traditional short-read sequencing and linear reference genome paradigm.

Long-read sequencing technologies have boosted our power in resolving complex regions of the genome, detecting structural variants, and assembling complete chromosomes [11–13]. In the recent years, these technologies have advanced in accuracy and yield, enabling variant detection at a large scale, with recent studies showcasing their potential in population-scale analyses [13,14].

Significant steps have also been taken to address the issue of the low ancestral diversity in the human genome reference. The Human Pangenome Reference Consortium has been working on constructing a reference that encompasses diverse genomes, and they have published the first draft of the pangenome, consisting of 47 genome assemblies from a diverse cohort [15]. However, there are still challenges to overcome in growing and refining this reference, both in sampling and in computational analysis. The current sampling efforts do not include Indigenous American or Aboriginal peoples (https://humanpangenome.org/samples/). Indigenous American people are genetically distinct compared to other populations in the United States, with a significant variation within the group itself [16]. Principal component analysis from a previous study including three ancestral populations from the 1000 Genomes database (European, East Asian, and African ancestries) and our Indigenous American population from Arizona (IAZ) showed a distinct cluster for IAZ on the principal component space [17]. Therefore, exploratory genomics studies using novel methods are warrented in indigenous populations to uncover previously unrecognized genetic variations and capture novel disease associations.

In this study, we aimed to improve identification of genetic variations that have not been previously studied among IAZ population. We utilized *de novo* genome assemblies to identify non-reference segments (NRSs) in this cohort. These NRSs represent novel genomic segments not present in the existing reference map. By analyzing these segments, both directly and by incorporating them into our whole-genome sequencing (WGS) variant calling pipeline, we sought to uncover previously unrecognized polymorphisms which can be assessed for a potential role in metabolic disease enriched in this underrepresented ethnic group.

## RESULTS

### *De novo* IAZ genome assemblies

DNA samples from one female (sample 1) and one male (Sample 2) Indigenous American from Arizona (IAZ) were sequenced using the PacBio HiFi SMRT sequencing platform. Summary reports for the raw PacBio sequencing data are shown in S1 Table. For both samples, the SMRT sequencing reactions produced approximately 64 Gb of high-fidelity subreads (>Q20) with a mean length of ∼12 kb, which were used to generate the *de novo* genome assemblies. Primary contigs with high contiguity were generated using HiFiasm assembler (Table 1). Assembly 1 (female IAZ) and assembly 2 (male IAZ) yielded a total genome length of 3.05 and 3.07 Gb, respectively, indicating high assembly quality regarding genome completeness when compared to the 2.96 Gb GRCh38/hg38 reference genome excluding gaps. Subsequent rounds of MUMmer alignment identified 19 Mb and 17 Mb of DNA sequence for assembly 1 and assembly 2, respectively, that could not be aligned to the hg38 reference genome. These non-reference sequences (NRSs) have a minimum length of 100 bps and a hg38 sequence identity < 80%.

**Table 1.**
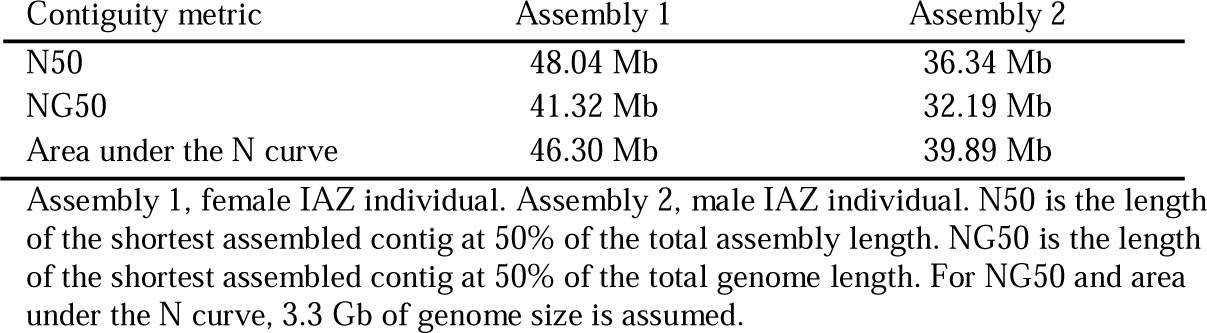
Summary statistics for the IAZ *de novo* assemblies.

### NRS size distribution for the IAZ *de novo* assemblies

The two IAZ *de novo* assemblies had similar numbers of NRSs and size distribution patterns with a total length of 19.3 Mb for assembly 1 (n = 2,177 NRSs) and 16.34 Mb for assembly 2 (n = 1908 NRSs) (Fig 1A-F). For both assemblies, most of the NRSs are 10 kb and shorter but only accounted for 14.4% and 13.4% of the total lengths for assembly 1 and assembly 2 respectively (Fig 1A and D). There were 130 NRSs for both assemblies with sizes greater than 10 kb–100 kb with medium sizes of 20.5 kb for assembly 1 and 22.7 kb for assembly 2 (Fig 1B and E). 93 NRSs are larger than 100 kb (assembly 1, n = 50; assembly 2 n = 43), including 3 outlier NRSs larger than 500 kb (Fig 1C and F).

**Fig 1:**
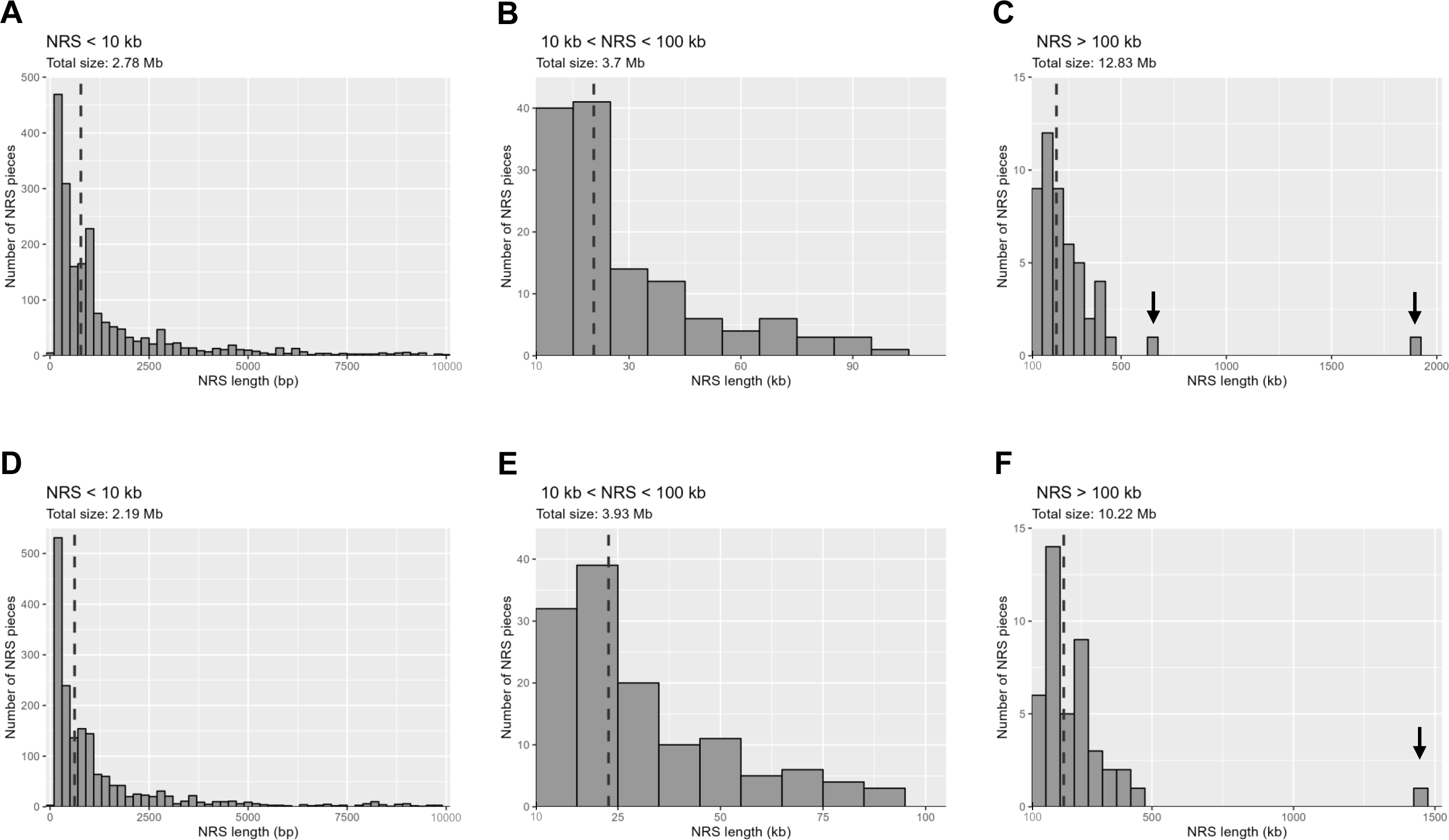
Histograms depicting the length distribution of NRSs identified in the indigenous Sample 1 (panels A to C) and Sample 2 (panels D to F). Dashed lines represent the median values for each histogram. **A**. Small NRSs ranging from 100 bp and 10 kb, with a median length of 776 bp. **B**. NRSs sized between 10 kb and 100 kb, with a median size of 20.5 kb. **C**. NRSs longer than 100 kb are shown, with a median size of 192 kb. Except for two outliers indicated by arrows, all NRSs in this group are shorter than 500 kb. The arrows in this panel highlight a 637 kb NRS and a 1.88 Mb NRS, both of which consist of simple repeats and microsatellites, and do not align to any hg38 contigs with high sequence identity. **D**. Small NRSs with varying lengths between 100 bp and 10 kb and with a median length of 617 bp. **E**. NRSs sized between 10 kb and 100 kb. The median size is 22.7 kb. **F**. NRSs longer than 100 kb, with a median size of 187.8 kb. There is a single NRS, marked with an arrow, exceeding 500 kb in length. This 1.47 Mb NRS comprises solely of simple repeats and microsatellites, and does not anchor to any hg38 contigs.

### Identification of repeat and transposable elements

Screening the NRSs for repeat and transposable elements using RepeatMasker revealed that approximately 95% of the NRSs in both assemblies primarily consisted of satellites, simple repeats, and a small fraction of LINE and SINE transposable elements (S1 Fig). The three largest NRSs with lengths of 637 kb, 1.88 Mb (assembly 1), and 1.47 Mb (assembly 2) were individually analyzed for their repeat content. They consisted almost entirely of satellites and simple repeats (ranging from 99.7% to 99.9%) rendering their alignment to chromosomes challenging.

### Mapping the NRSs to the GRCh38/hg38 reference genome

NRSs totaling 241 kb could be placed into 40 distinct locations within the primary scaffold of GRCh38/hg38. Only 4 of the NRSs totaling 7.6 kb were placed into centromeric regions suggesting that there was no enrichment of centromeric DNA which is in contrast to long-read sequencing projects involving other populations [18, 19]. However, when the alignments for all of the NRSs with at least one anchored flanking site were analyzed, the percentage of NRSs aligning to centromeric or telomeric regions was ∼30%. The 233.4 kb non-centromeric NRSs includes a large 215 kb insertion aligning to chrY and ∼18.4 kb of NRS anchoring to 13 autosomal chromosomes, chrX, and 2 pseudoautosomal regions on chrY. Among the 36 non-centromeric NRSs, 17 are located within intronic regions of 13 protein coding and 4 non-protein-coding genes. Three NRSs were present in both assemblies and only 1, *HCN2*-NRS, was identical in size and therefore was selected for further characterization (Table 2). The *HCN2*-NRS in not present in the reference genomes, including the T2T-CHM13 map, suggesting that the *HCN2*-NRS may not exist in the DNA samples used for reference genome construction. *HCN2*-NRS is located 79 bp downstream of exon 3 and replaces 228 bp of the reference sequence including a small portion of a predicted cis-regulatory element (CRE). Individuals who carry the 187 bp HCN2-NRS also have an adjacent 100 bp deletion (S2 Fig). The *HCN2*-NRS matches (100% identity) with 187 bps downstream of exon 4 spanning a putative CRE containing various transcription factor binding sites suggesting that *HCN2*-NRS may have functional importance (Fig 2, S2 Fig).

**Fig 2.**
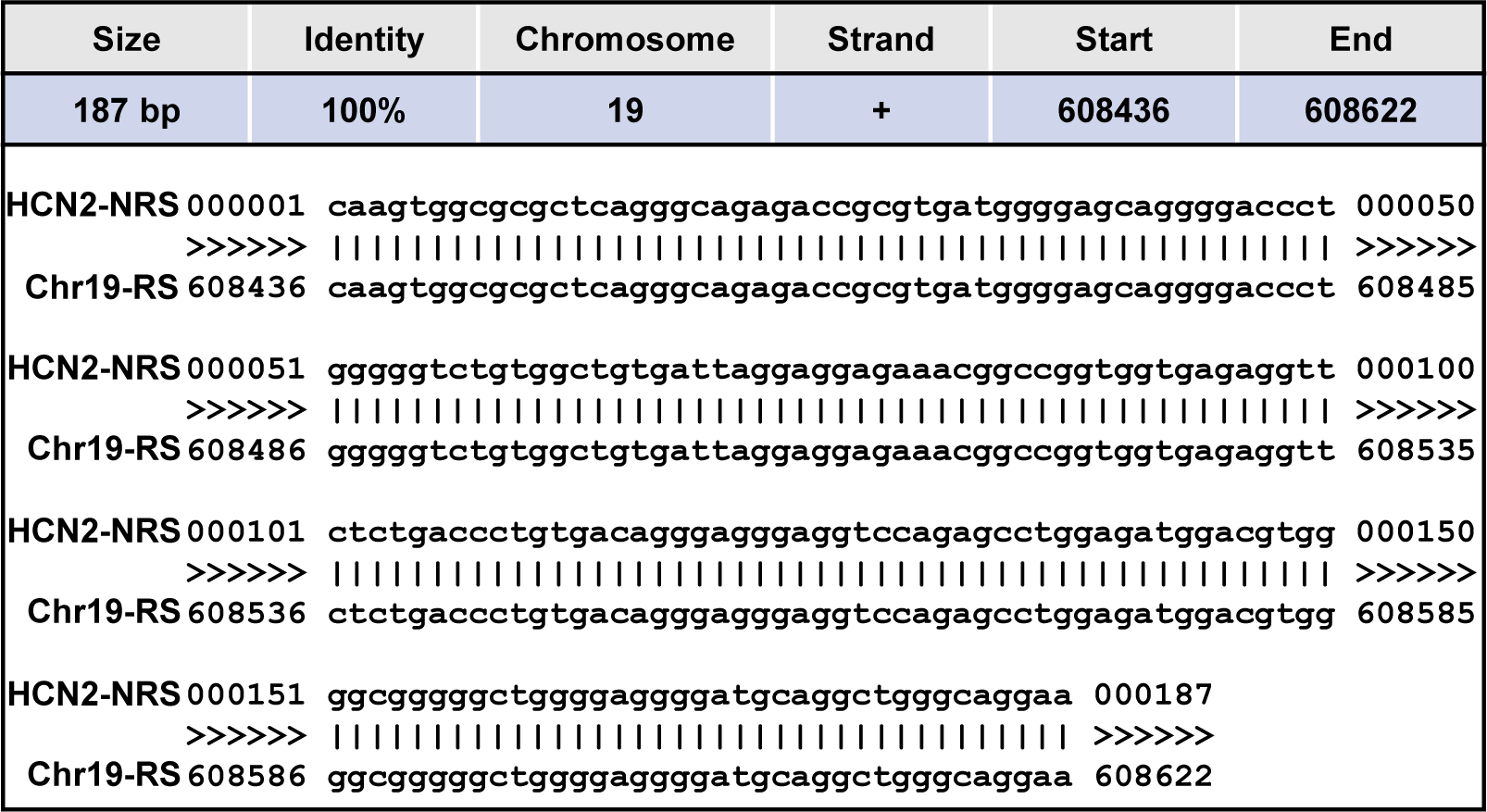
HCN2-NRS and chr19-RS side by side alignment. Source: UCSC Genome Browser (GRCh38/hg38).

**Table 2.** Non-reference sequences anchoring within genes.

| Gene | Insertion region<sup>a</sup> | NRS size (bps) | Both assemblies |
| --- | --- | --- | --- |
| *LINC02270* | Chr4:12237666–12237666 | 139 | No |
| *F11-AS1* | Chr4:186435038–186435150 | 4714 | No |
| *BTNL9* | Chr5:181046016–181046016 | 386 | No |
| *DYNC2I1* | Chr7:158915724–158915724 | 679 | No |
| *MROH5* | Chr8:141492602–141492602 | 1722 | No |
| *GTPBP4* | Chr10:1011004–1011004 | 341 | No |
| *ART1* | Chr11:3653879–3653937 | 877 | Yes<sup>b</sup> |
| *PDGFD* | Chr11:104078565–104078565 | 579 | No |
| *LOC728715* | Chr12:9404585–9404585 | 836 | Yes<sup>b</sup> |
| *GNPTAB* | Chr12:101747637–101747644 | 658 | No |
| *MPG* | Chr16:81854–81857 | 344 | No |
| *CACNA1H* | Chr16:1186044–1186087 | 466 | No |
| *HCN2* | Chr19:605301–605528 | 187 | Yes |
| *ADGRG2* | ChrX:19074564–19074564 | 429 | No |
| *RGN* | ChrX:47089179–47089179 | 1127 | No |
| *CRLF2* | ChrY:1198256–1198271<sup>c</sup> | 287 | No |
| *P2RY8* | ChrY:1496850–1496850<sup>c</sup> | 136 | No |

NRS, non-reference sequence. <sup>a</sup>Position based on GRCh38/hg38. Indicated intervals are replaced by NRSs.
<sup>b</sup>NRS sizes slightly different between the two assemblies. <sup>c</sup>Pseudoautosomal region.

### Population frequency and association analyses for *HCN2*-NRS

To confirm the presence of the *HCN2*-NRS, the region encompassing the putative NRS was PCR amplified and sequenced in the two IAZ individuals used for the *de novo* genome assembly along with some of their family members. Of the 10 individuals sequenced, 1 was heterozygous and 9 were homozygous for *HCN2*-NRS suggesting that the NRS segment was common in the IAZ population. To determine the frequency of *HCN2*-NRS in IAZ, ∼3,400 IAZ individuals were genotyped using a custom designed TaqMan probe that detects the presence of the NRS. The *HCN2*-NRS is the minor allele with a frequency of 0.45. Genotyping was repeated for several individuals by PCR to verify that the custom probe was working properly (S3 Fig). For comparison, the *HCN2*-NRS was also genotyped in the African Ancestry in Southwest USA (ASW) and British from England and Scotland (GBR) DNA panels from the 1000 genomes project. *HCN2*-NRS was not as common in these two ethnic groups with frequencies of 0.03 and 0.15 in the ASW and GBR populations, respectively. Despite being enriched in IAZ, the *HCN2*-NRS allele was not significantly associated with any obesity or type 2 diabetes-related traits which are highly prevalent in this population (data not shown).

### Structural variation in the *de novo* assemblies

Structural variants (SV) were analyzed directly from the *de novo* assemblies using Assemblytics, which distinguishes between insertions/deletions and contractions/expansions of repeat elements [20]. A total of 12,265 and 12,133 SVs were detected for Sample 1 and Sample 2, respectively, which is similar to an Assemblytics report for a human reference assembly (11,206 SVs) [20] but lower than the number of SVs reported in other studies that used assembly-based or long-read mapping approaches [18, 21]. The lower number of SVs in our study can be attributed to the stringency of our SV detection method. For both samples, the larger SVs (500bp–10,000bp) are predominately composed of tandem expansions (S4 Fig). The distribution graphs also show peaks of SV counts at around 300 bp and 6,000 bp (S4 Fig), a size distribution pattern similar to a previous report [18].

### Variant calling using the amended hg38-NRS map

Variant calling using our short-read WGS data generated for 387 IAZ individuals and the amended hg38-NRS map identified 11,130,171 variants. Using the new hg38-NRS map identified 161,502 gained single nucleotide variants (SNVs) including 695 missense substitutions that were not captured when variant calling was performed using the hg38 reference map (Fig 3). Additionally, 180,940 SNVs that were captured using the hg38 map were not detected (lost) with the hg38-NRS map (Fig 3). Most of the gained SNVs (n = 111,976) were low-frequency variants; while the remaining gained SNVs (n = 49,526) were detected in ≥ 5% of the WGS samples. The accuracy of our variant calling with the hg38-NRS map was assessed by sequencing 25 gained SNVs (15 reported and 10 not reported in dbSNP). For the 15 dbSNPs, 13 of the gained SNVs were verified and for the 10 non-dbSNPs, 6 were validated. Most of the non-dbSNPs SNVs reside in repetitive regions of the genome which may account for the reduced accuracy.

**Fig 3.**
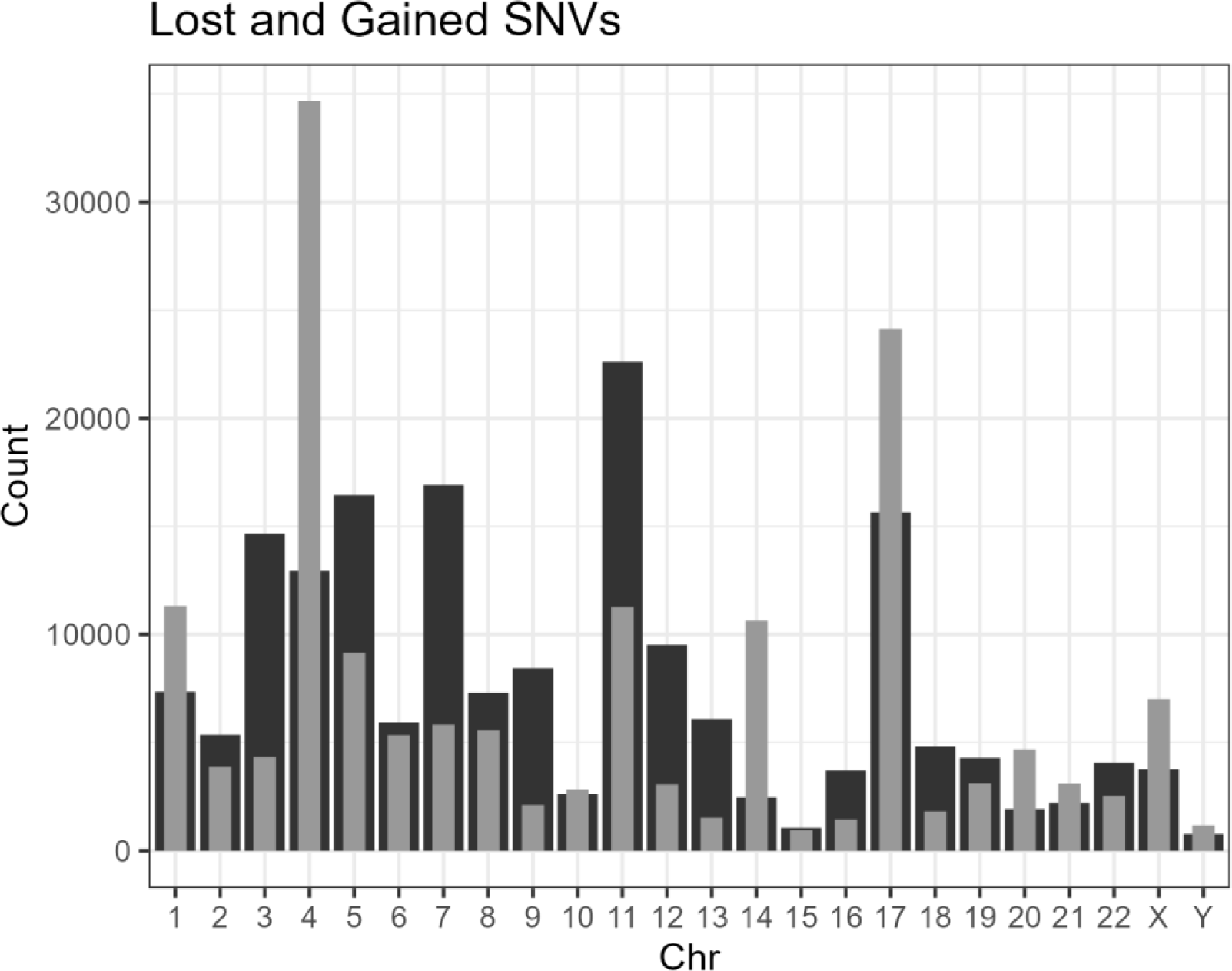
Number of SNVs gained (light gray) and lost (dark gray) per chromosome for the cohort of 387 short-read WGS samples when analyzed with the hg38-NRS map compared to hg38 map.

## DISCUSSION

In this study, our goal was to enhance the identification of genetic variations that remain largely unexplored within an Indigenous American cohort from Arizona. We focused on identifying NRSs in *de novo* genome assemblies constructed for the two participants from the IAZ cohort. The newly identified NRSs signify novel genomic regions not found in the current GRCh38/hg38 reference map. By incorporating these NRS segments into our WGS variant calling pipeline, we aimed to uncover previously undetected polymorphisms.

The *de novo* assemblies were constructed using long-read PacBio HiFi sequencing which produces highly accurate long-read (>10 kb) sequencing data [22]. The sizes for the *de novo* genome assemblies were similar to hg38 and exhibited high contiguity with a mean N50 value of 42.2 Mb. We identified > 10,000 SVs per *de novo* genome assembly which is in line with the expectations for long-read sequencing [23]. This assembly-based approach for SV detection proved particularly advantageous for capturing variants of substantial size [24], and the categorization facilitated by Assemblytics allowed us to differentiate between SVs occurring in repetitive regions, such as tandem expansions and repeat contractions, and those in non-repetitive regions (insertions/deletions). It is worth noting that the largest insertions and deletions (> 500 kb) often resided in regions characterized by high repeat content. We discovered 220 shared deletions (> 500 kb) in both *de novo* assemblies with 90 SVs located within genes (data not shown). The application of a high-throughput genotyping approach with custom CNV probes could potentially reveal commonly occurring deletions and uncover potential associations with diseases. However, probe designs for these regions, predominantly repetitive in content, would pose a challenge.

The detected NRSs in two assemblies were found to be highly enriched in repeat sequences, especially satellites and simple repeats. This high repeat content was consistent with the previous literature on non-reference genomic parts [21, 25], and rendered the precise placement of NRSs into chromosomes challenging. We identified 1.5 Mb of non-repetitive NRS per assembly and 241 kb was anchored to specific chromosomal regions in the primary assembly of hg38. The largest NRS (215 kb) was anchored to chrY. This large NRS is probably a result of chrY being incomplete in the current hg38 map due to its complex repeat structure [26].

Several of the NRSs anchored to intronic regions and we focused on a particular NRS in *HCN2*. *HCN2*-NRS was detected in both de novo assemblies and genotyping showed that the *HCN2*-NRS allele is common (MAF = 0.45) in the IAZ population. We also determined that *HCN2*-NRS is present in Caucasians and African Americans but at lower frequencies (MAF = 0.15 and 0.03, respectively).

The *HCN2*-NRS matches (100% identity) with a region located downstream of exon 4 that encompasses a putative *cis*-regulatory element suggesting that individuals carrying the *HCN2*-NRS may have additional transcription factor binding sites which may potentially influence gene expression. *HCN2* encodes for a voltage-gated cation channel predominantly expressed in the heart and the brain and plays a crucial role in migraine, inflammatory, and neuropathic pain including diabetic neuropathy [27–29]. HCN2 ion channels initiate neuropathic pain by modulating action potential firing in nociceptor neurons involved in sensing pain intensity [30]. Compared to Caucasians, chronic widespread and localized musculoskeletal pain disorders are uncommon among IAZ [31], however, IAZ have high rates of diabetic neuropathy and pain from rheumatoid arthritis [32, 33]. It is possible that the common *HCN2*-NRS allele might modestly contribute to diabetic neuropathy since there are multiple rare intronic variants with nominal associations with diabetic neuropathy reported in the T2D Knowledge Portal (https://t2d.hugeamp.org/). However, the current study has no data on diabetic neuropathy pain to address this hypothesis.

Our validation assessment for “gained” variants, namely the subset of genomic variants that were only called using hg38-NRS as the reference, using Sanger sequencing underlines the reliability of our WGS data analysis with the extended reference map. However, the calls arising from repetitive regions provided lower rates of true positives. Further research is needed to understand the functional significance of the gained variants and potential contributions to diseases among IAZ population.

The overall findings we presented have expanded our understanding of genetic diversity within Indigenous American populations. The presence of unique genetic elements and variants unrepresented in existing reference maps demonstrates the importance of delving into the genomics of underrepresented populations. By combining long-read sequencing and novel NRS identification methods, we were able to uncover previously elusive genetic variations. This study not only highlights the value of studying isolated populations but also emphasizes the need for diverse and comprehensive reference maps to accurately capture the complexity of human genetic diversity. Our work contributes to the broader goal of enhancing genomic research’s inclusivity and represents a step toward more precise understanding of genetic factors underlying health disparities in underrepresented populations.

## METHODS

### Long-read sequencing and *de novo* assembly

Two DNA samples isolated from one female and one male individual were sequenced using the PacBio HiFi SMRT sequencing platform at the DNA Sequencing Center at Brigham Young University (Provo, UT). SMRT bell adapted DNA libraries were size selected and ∼15 kb fractions were run on 3 SMRT cells (8M trays) for a duration of 30 hours for each sample. Bioinformatic analysis for the construction of the IAZ *de novo* assemblies was done by the DNA Sequencing Center at Brigham Young University. Selected quality reads were assembled using Hifiasm version 0.11 (r302). Continuity of the reads and other summary statistics were calculated using the caln50 package (https://github.com/lh3/calN50).

### Detection of non-reference sequences and structural variants in the *de novo* assemblies

All computational analyses were run on the high-performance computing environment NIH HPC Biowulf cluster (http://hpc.nih.gov). Visualization was performed on RStudio using ggplot2 package of R (version 4.3.1).

The *de novo* assemblies were compared to the Genome Reference Consortium Human Build 38 patch release 13 (GRCh38.p13/hg38) using the alignment script of NUCmer from the MUMmer package (version 4.0.0beta2). Each assembly was aligned to GRCh38.p13/hg38 with the parameters--maxmatch −l 150-c 400. The resulting alignments were filtered to retain only the single best alignment for each contig using the delta-filter -q command. The show-coords tool was then used to extract the coordinates and percent identity of the aligned contigs. To identify the contigs that failed to align to hg38 (< 80% sequence identity), a custom script (available at https://github.com/cigdemkoroglu/NRS/blob/main/unaligned_contigs.R) was used to parse the output of the alignment and extract the unaligned contigs and contigs with low identity. Bedtools version 2.29.2 was employed to generate fasta files containing only the unaligned portions of the assemblies (NRSs). To improve the alignment sensitivity and capture more contigs, the alignment workflow was repeated on the initial output of NRSs with relaxed NUCmer settings (--maxmatch −l 100-c 200). This allowed for a more permissive alignment approach to increase the number of contigs that could be aligned to the reference. Final NRSs were analyzed for repetitive sequences using RepeatMasker and Assemblytics was used to detect structural variants > 50 bp (http://assemblytics.com/).

### Anchoring the NRSs to the GRCh38.p13/hg38 primary assembly

For each NRS, 100 bp of flanking sequences either side of the NRSs were aligned to the nucleotide collection database using the blastn. Contigs from the *de novo* assemblies with both flanking sites anchoring to the primary assembly chromosomes of hg38 were selected and appended to hg38 to generate an extended reference genome (hg38-NRS) using perEditor (https://systemsbio.ucsd.edu/perEditor/). NRSs spanning a reference region longer than the NRS itself and those aligning to the same region, indicating repeat expansions, were not incorporated into the hg38-NRS reference genome. In cases where NRSs overlapped between both assemblies, only the region from assembly 1 was included.

### Variant calling using hg38 and hg38-NRS reference genomes

Whole genomes for 387 IAZ individuals were previously sequenced using Illumina short-read chemistry (30× average coverage). Variant discovery was performed in parallel on whole genome sequencing data from 387 IAZ samples using either hg38 or hg38-NRS as the reference. GATK 4.2.6 was used following the best practices workflow, which includes base quality score recalibration, variant calling and joint genotyping by HaplotypeCaller and GenotypeGVCFs, and variant quality score recalibration. Databases for the known variation in the GATK resource bundle were position adjusted based on hg38-NRS using a custom lift-over script (available at https://github.com/cigdemkoroglu/NRS/blob/main/lift_database.R). Monomorphic variants, variants with a call rate <95%, variants with quality scores below the truth sensitivity level 99% threshold, and variants with discordant genotypes between the duplicate pairs were removed from the final vcf files using VCFtools.

### Identification of gained single nucleotide variants (SNVs) in the hg38-NRS reference genome

To compare variant calling between hg38 and hg38-NRS reference genomes, ANNOVAR was first used to annotate the variants identified with hg38. Next, to annotate the variants identified with hg38-NRS and to ensure compatibility between the two reference maps, the hg38-NRS variant dataset was lifted-over using custom scripts (https://github.com/cigdemkoroglu/NRS). Lastly, to identify lost and gained SNVs, the hg38 and hg38-NRS annotated datasets were compared using the LostGained tool (https://github.com/ cigdemkoroglu/NRS/blob/main /LostGained.R). Among the gained variants, 15 missense variants that are reported in dbSNP but were not detected by our in-house genotyping studies using the Affymetrix SNP Array 6.0 or our whole-genome and whole-exome sequencing studies along with 10 randomly selected novel SNVs were verified by sequencing.

### Genotyping of the *HCN2*-NRS

Genotyping of the *HCN2*-NRS in the Indigenous Americans from Arizona (IAZ) was performed using DNA isolated from 3,410 individuals who participated in a community-based longitudinal study examining type 2 diabetes-related traits conducted by the National Institute of Diabetes and Digestive and Kidney Diseases (NIDDK) in Phoenix, Arizona between the years 1965 and 2007 [34]. All participants in the study were self-reported full-heritage Indigenous American (all eight great grandparents identified as IAZ). The *HCN2*-NRS was also genotyped in the African Ancestry in Southwest USA (ASW) and British from England and Scotland (GBR) DNA panels from the 1000 genomes project. The ASW and GBR DNA samples were obtained from the NHGRI Sample Repository for Human Genetic Research at the Coriell Institute for Medical Research: Repository IDs MGP00015 and MGP00003, respectively. Genotyping of the *HCN2*-NRS was done using custom designed TaqMan probes that detected the presence of either the reference sequence (RS) or NRS. (Thermo Fisher Scientific, Waltham, MA). Statistical analyses were performed using the software of the SAS Institute (Cary, NC, USA) as previously described [35].

## Acknowledgments

We thank Chris Wiedrich for his help in genotype validation, Sayuko Kobes for running association tests, and Paolo Piaggi for providing statistical consultation. This work was supported by the Intramural Research Program of the NIDDK, NIH. We utilized the computational resources of the NIH HPC Biowulf cluster (http://hpc.nih.gov).

## Author contributions

Conceptualization: Cigdem Koroglu, Clifton Bogardus Study design & data analysis: Cigdem Koroglu Consultation regarding data analysis: Peng Chen Supervision: Leslie J. Baier Software: Serdar Altok, Peng Chen, Cigdem Koroglu Computational troubleshooting: Cigdem Koroglu, Serdar Altok Genotyping: Michael Traurig, Cigdem Koroglu Visualization: Cigdem Koroglu, Michael Traurig Writing-original draft: Cigdem Koroglu Writing-review & editing: Leslie J. Baier, Cigdem Koroglu, Michael Traurig, Peng Chen, Clifton Bogardus, Serdar Altok

## Competing interests

The authors declare no competing interests.

## Notes

https://drive.google.com/drive/folders/19y4RUmsRDvm6_PJnAIdGyztBcZ44VAlI?usp=sharing

### Competing Interest Statement

The authors have declared no competing interest.

### Summary of Updates

Supplementary files are added

## References

1. Gurdasani D, Barroso I, Zeggini E, Sandhu MS. Genomics of disease risk in globally diverse populations. Nat Rev Genet. 2019; 20(9): 520–535.

2. Manrai AK, Funke BH, Rehm HL, Olesen MS, Maron BA, Szolovits P, et al. Genetic Misdiagnoses and the Potential for Health Disparities. N Engl J Med. 2016; 375(7): 655–665.

3. Popejoy AB, Fullerton SM. Genomics is failing on diversity. Nature. 2016; 538(7624): 161-164.

4. All of Us Research Program Investigators; Denny JC, Rutter JL, Goldstein DB, Philippakis A, Smoller JW, et al. The “All of Us” Research Program. N Engl J Med. 2019; 381(7): 668-676.

5. GenomeAsia100K Consortium. The GenomeAsia 100K Project enables genetic discoveries across Asia. Nature. 2019; 576(7785): 106-111.

6. Mulder N, Abimiku A, Adebamowo SN, de Vries J, Matimba A, Olowoyo P, et al. H3Africa: current perspectives. Pharmgenomics Pers Med. 2018; 11: 59–66.

7. Chheda H, Palta P, Pirinen M, McCarthy S, Walter K, Koskinen S, et al. Whole-genome view of the consequences of a population bottleneck using 2926 genome sequences from Finland and United Kingdom. Eur J Hum Genet. 2017; 25(4): 477–484.

8. Beyter D, Ingimundardottir H, Oddsson A, Eggertsson HP, Bjornsson E, Jonsson H, et al. Long-read sequencing of 3,622 Icelanders provides insight into the role of structural variants in human diseases and other traits. Nat Genet. 2021; 53(6): 779–786.

9. Wu Z, Jiang Z, Li T, Xie C, Zhao L, Yang J, et al. Structural variants in the Chinese population and their impact on phenotypes, diseases and population adaptation. Nat Commun. 2021; 12(1): 6501.

10. Schneider VA, Graves-Lindsay T, Howe K, Bouk N, Chen HC, Kitts PA, et al. Evaluation of GRCh38 and de novo haploid genome assemblies demonstrates the enduring quality of the reference assembly. Genome Res. 2017; 27(5): 849–864.

11. Chaisson MJ, Huddleston J, Dennis MY, Sudmant PH, Malig M, Hormozdiari F, et al. Resolving the complexity of the human genome using single-molecule sequencing. Nature. 2015; 517(7536): 608-611.

12. Logsdon GA, Vollger MR, Eichler EE. Long-read human genome sequencing and its applications. Nat Rev Genet. 2020; 21(10): 597–614.

13. Marx V. Method of the year: long-read sequencing. Nat Methods. 2023; 20(1): 6–11.

14. De Coster W, Weissensteiner MH, Sedlazeck FJ. Towards population-scale long-read sequencing. Nat Rev Genet. 2021; 22(9): 572–587.

15. Liao WW, Asri M, Ebler J, Doerr D, Haukness M, Hickey G, et al. A draft human pangenome reference. Nature. 2023; 617(7960): 312-324.

16. Redd AJ, Chamberlain VF, Kearney VF, Stover D, Karafet T, Calderon K, et al. Genetic structure among 38 populations from the United States based on 11 U.S. core Y chromosome STRs. J Forensic Sci. 2006; 51(3): 580–585.

17. Kim HI, Ye B, Gosalia N; Regeneron Genetics Center; Köroğlu Ç, Hanson RL, et al. Characterization of Exome Variants and Their Metabolic Impact in 6,716 American Indians from the Southwest US. Am J Hum Genet. 2020; 107(2): 251-264.

18. Ameur A, Che H, Martin M, Bunikis I, Dahlberg J, Höijer I, et al. De Novo Assembly of Two Swedish Genomes Reveals Missing Segments from the Human GRCh38 Reference and Improves Variant Calling of Population-Scale Sequencing Data. Genes (Basel). 2018; 9(10): 486.

19. Gao Y, Yang X, Chen H, Tan X, Yang Z, Deng L, et al. A pangenome reference of 36 Chinese populations. Nature. 2023; 619(7968): 112-121.

20. Nattestad M, Schatz MC. Assemblytics: a web analytics tool for the detection of variants from an assembly. Bioinformatics. 2016; 32(19): 3021–3023.

21. Bert P, Audano PA, Zhu Q, Rodriguez-Martin B, Porubsky D, Bonder MJ, et al. Haplotype-resolved diverse human genomes and integrated analysis of structural variation. Science. 2021; 372(6537): eabf7117.

22. Hon T, Mars K, Young G, Tsai YC, Karalius JW, Landolin JM, et al. Highly accurate long-read HiFi sequencing data for five complex genomes. Sci Data. 2020; 7(1): 399.

23. Ho SS, Urban AE, Mills RE. Structural variation in the sequencing era. Nat Rev Genet. 2020; 21(3): 171–189.

24. Mahmoud M, Gobet N, Cruz-Dávalos DI, Mounier N, Dessimoz C, Sedlazeck FJ. Structural variant calling: the long and the short of it. Genome Biol. 2019; 20(1): 246.

25. Li R, Tian X, Yang P, Fan Y, Li M, Zheng H, et al. Recovery of non-reference sequences missing from the human reference genome. BMC Genomics. 2019; 20(1): 746.

26. Skaletsky H, Kuroda-Kawaguchi T, Minx PJ, Cordum HS, Hillier L, Brown LG, et al. The male-specific region of the human Y chromosome is a mosaic of discrete sequence classes. Nature. 2003; 423(6942): 825-837.

27. Young GT, Emery EC, Mooney ER, Tsantoulas C, McNaughton PA. Inflammatory and neuropathic pain are rapidly suppressed by peripheral block of hyperpolarisation-activated cyclic nucleotide-gated ion channels. Pain. 2014; 155(9): 1708–1719.

28. Tsantoulas C, Ng A, Pinto L, Andreou AP, McNaughton PA. HCN2 Ion Channels Drive Pain in Rodent Models of Migraine. J Neurosci. 2022; 42(40): 7513–7529.

29. Tsantoulas C, Laínez S, Wong S, Mehta I, Vilar B, McNaughton PA. Hyperpolarization-activated cyclic nucleotide-gated 2 (HCN2) ion channels drive pain in mouse models of diabetic neuropathy. Sci Transl Med. 2017; 9(409): eaam6072.

30. Emery EC, Young GT, Berrocoso EM, Chen L, McNaughton PA. HCN2 ion channels play a central role in inflammatory and neuropathic pain. Science. 2011; 333(6048): 1462-1466.

31. Jacobsson LT, Nagi DK, Pillemer SR, Knowler WC, Hanson RL, Pettitt DJ, et al. Low prevalences of chronic widespread pain and shoulder disorders among the Pima Indians. J Rheumatol. 1996; 23(5): 907–909.

32. Jaiswal M, Fufaa GD, Martin CL, Pop-Busui R, Nelson RG, Feldman EL. Burden of Diabetic Peripheral Neuropathy in Pima Indians With Type 2 Diabetes. Diabetes Care. 2016; 39(4): e63–64.

33. Del Puente A, Knowler WC, Pettitt DJ, Bennett PH. High incidence and prevalence of rheumatoid arthritis in Pima Indians. Am J Epidemiol. 1989; 129(6): 1170–1178.

34. Knowler WC, Bennett PH, Hamman RF, Miller M. Diabetes incidence and prevalence in Pima Indians: a 19-fold greater incidence than in Rochester, Minnesota. Am J Epidemiol. 1978; 108(6): 497–505.

35. Day SE, Traurig M, Kumar P, Piaggi P, Koroglu C, Kobes S, et al. Functional variants in cytochrome b5 type A (CYB5A) are enriched in Southwest American Indian individuals and associate with obesity. Obesity (Silver Spring). 2022; 30(2): 546–552.

