## Supplementary material for "*De novo* genome assemblies from two Indigenous Americans from Arizona identify new polymorphisms in non-reference sequences": S1 Table 1

| **S1 Table**. Summary reports for the raw PacBio sequencing data. | | | |
| --- | --- | --- | --- |
| Sample 1  (Female IAZ) | SMRT cell 1 | SMRT cell 2 | SMRT cell 3 |
| Polymerase Read Bases | 280,457,580,897 | 399,427,202,940 | 406,977,340,901 |
| Polymerase Reads | 3,199,750 | 4,347,153 | 5,086,721 |
| Polymerase Read Length (mean) | 87,649 | 91,882 | 80,007 |
| Polymerase Read N50 | 187,203 | 190,600 | 165,676 |
| Subread Length (mean) | 12,759 | 11,103 | 11,361 |
| Subread N50 | 14,457 | 11,887 | 12,018 |
| Insert Length (mean) | 14,693 | 13,934 | 15,651 |
| Insert N50 | 15,562 | 13,036 | 14,106 |
| Unique Molecular Yield | 44,588,875,776 | 56,367,218,688 | 73,775,841,280 |
| Sample 2  (Male IAZ) | SMRT cell 1 | SMRT cell 2 | SMRT cell 3 |
| Polymerase Read Bases | 157,284,829,272 | 221,684,966,545 | 383,044,553,659 |
| Polymerase Reads | 1,704,132 | 2,574,815 | 4,356,450 |
| Polymerase Read Length (mean) | 92,296 | 86,097 | 87,925 |
| Polymerase Read N50 | 187,804 | 182,428 | 182,369 |
| Subread Length (mean) | 13,537 | 10,913 | 11,110 |
| Subread N50 | 14,662 | 11,765 | 11,712 |
| Insert Length (mean) | 16,117 | 12,973 | 14,887 |
| Insert N50 | 16,214 | 12,723 | 13,266 |
| Unique Molecular Yield | 26,025,361,408 | 31,336,163,328 | 59,633,332,224 |
| Subread N50 | 14,662 | 11,765 | 11,712 |

IAZ, Indigenous American from Arizona.
