## Supplementary material for "*De novo* genome assemblies from two Indigenous Americans from Arizona identify new polymorphisms in non-reference sequences": S1-S4 Figs

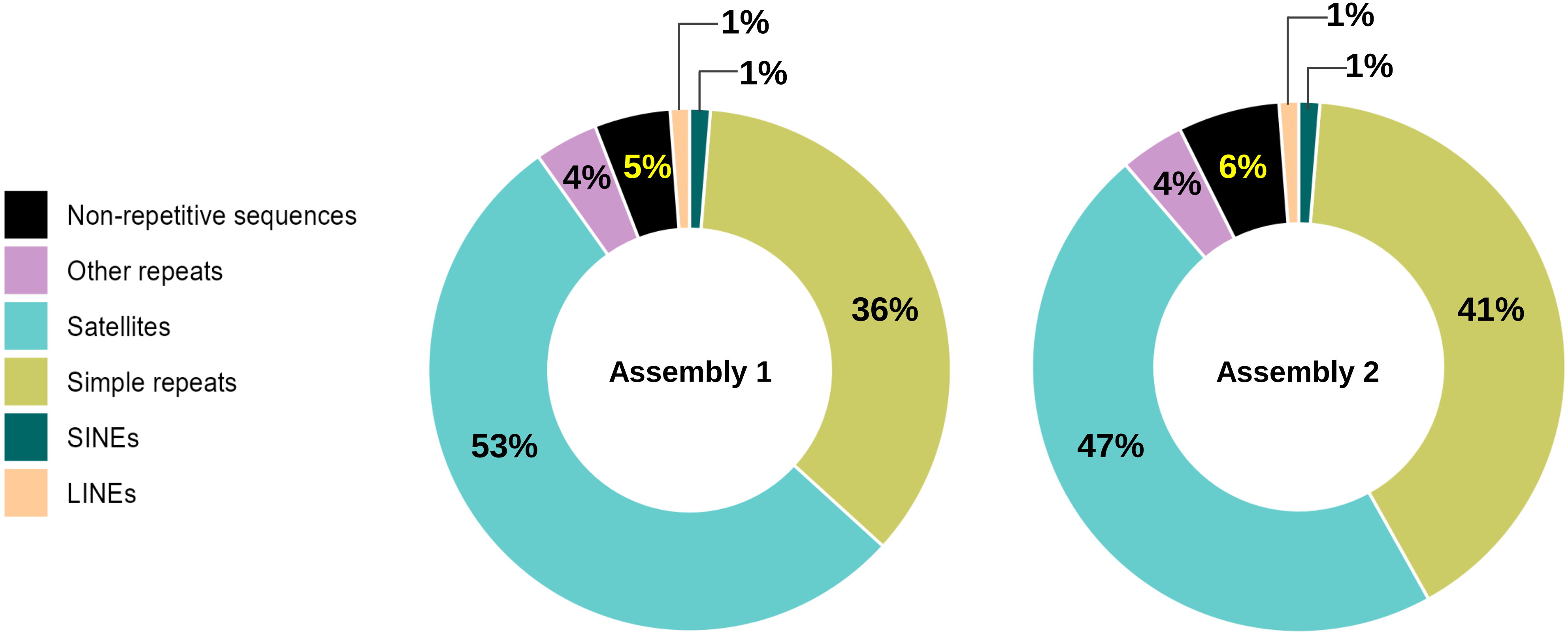


**S1 Figure.** Repeat content for the NRSs detected in both *de novo* assemblies.

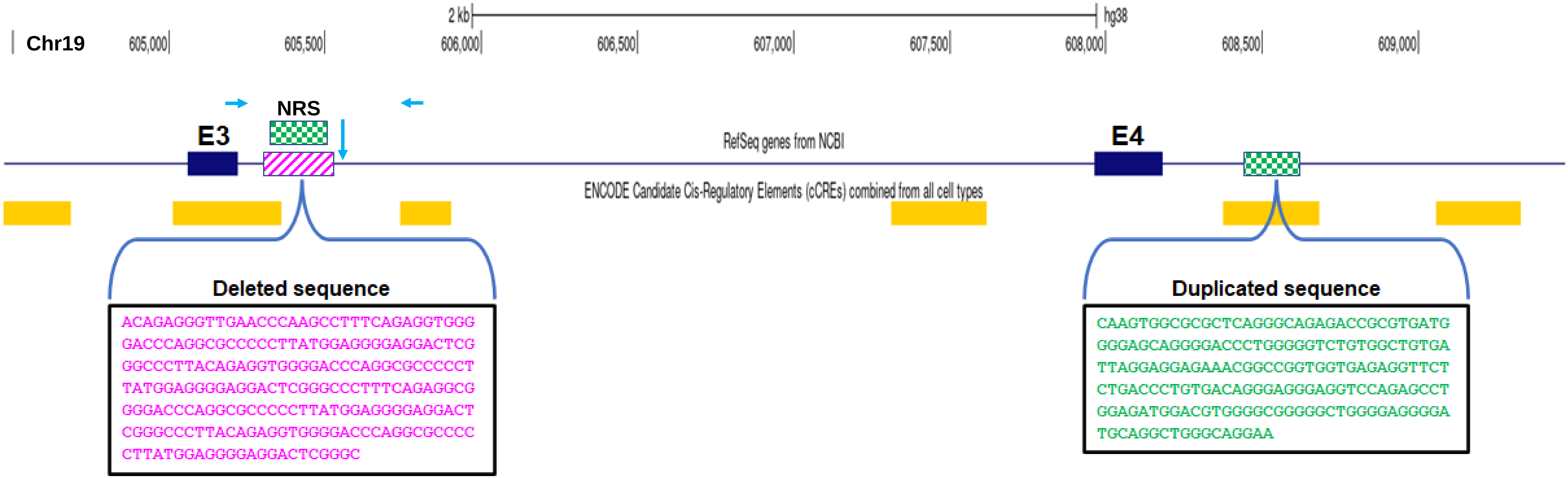


**S2 Figure.** Schematic showing the location of the 187 bp *HCN2*-NRS adjacent to exon 3. The NRS is identical to a 187 bp region located in a predicted *cis*-regulatory element downstream of exon 4. Light blue horizontal arrows indicate the locations of the primers used to screen for the presence of the 187 bp NRS. Light blue vertical arrow indicates the approximate start site of an additional 104 bp deletion present in individuals who carry the 187 bp NRS. The figure was adapted from the UCSC Genome Browser on Human (GRCh38/hg38).

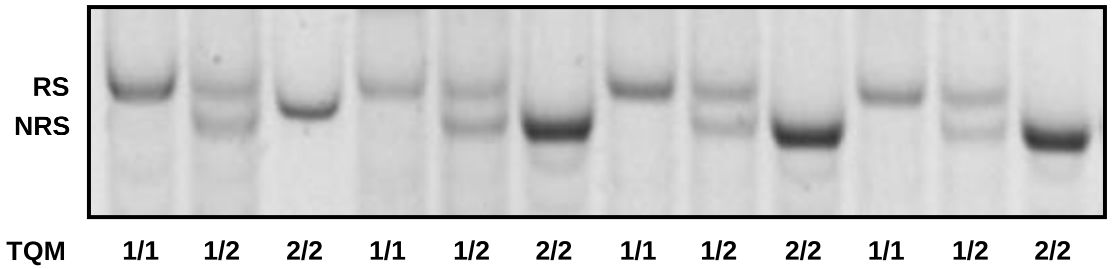


**S3 Figure.** Representative gel showing the comparison between the *HCN2*-NRS genotyping by PCR and custom TaqMan probe. RS, reference sequence. NRS, non-reference sequence. TQM, TaqMan probe genotypes. 1/1, homozygous reference sequence. 1/2, heterozygous. 2/2, homozygous for the non-reference sequence.

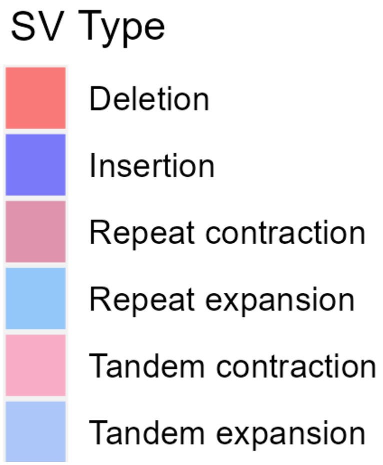


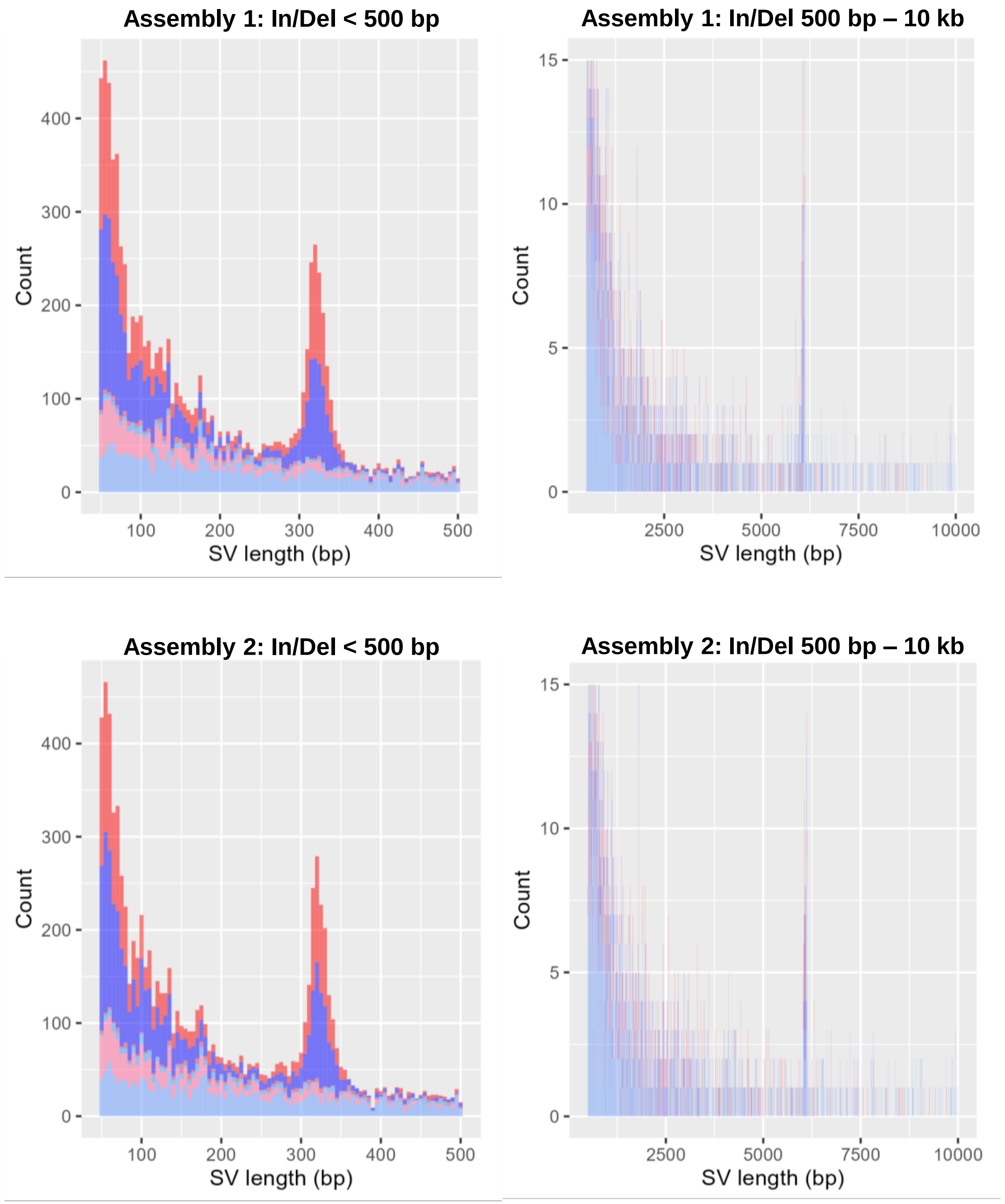


**S4 Figure.** The size distribution of structural variants detected in *de novo* assembly 1 (top panels) and *de novo* assembly 2 (bottom panels). Histograms show the lengths of insertions (insertion, repeat expansion, tandem expansion; shades of blue) and deletions (deletion, repeat contraction, tandem contraction; shades of red).
